# Chemogenetic silencing of hippocampus and amygdala reveals a double dissociation in periadolescent obesogenic diet-induced memory alterations

**DOI:** 10.1101/2020.07.20.212258

**Authors:** Fabien Naneix, Ioannis Bakoyiannis, Marianela Santoyo-Zedillo, Clémentine Bosch-Bouju, Gustavo Pacheco-Lopez, Etienne Coutureau, Guillaume Ferreira, OBETEEN Consortium

**Author notes:** FN and IB contributed equally to this work. EC and GF jointly directed this work.

## Abstract

In addition to numerous metabolic comorbidities, obesity is associated with several adverse neurobiological outcomes, especially learning and memory alterations. Obesity prevalence is rising dramatically in youth and is persisting in adulthood. This is especially worrying since adolescence is a crucial period for the maturation of certain brain regions playing a central role in memory processes such as the hippocampus and the amygdala. We previously showed that periadolescent exposure to obesogenic high-fat diet (HFD) had opposite effects on hippocampus- and amygdala-dependent memory, impairing the former and enhancing the latter. However, the causal role of these two brain regions in periadolescent HFD-induced memory alterations remains unclear. Here, we first showed that periadolescent HFD induced long-term, but not short-term, object recognition memory deficits, specifically when rats were exposed to a novel context. Using chemogenetic approaches to inhibit targeted brain regions, we then demonstrated that recognition memory deficits are dependent on the activity of the ventral hippocampus, but not the basolateral amygdala. On the contrary, the HFD-induced enhancement of conditioned odor aversion requires specifically amygdala activity. Taken together, these findings suggest that HFD consumption throughout adolescence impairs long-term object recognition memory through the overactivation of the ventral hippocampus during memory acquisition. Moreover, these results further highlight the bidirectional effects of adolescent HFD on hippocampal and amygdala functions.

## 1 INTRODUCTION

Obesity is one of the most important public health challenges and is linked to the overconsumption of energy-dense food combined with a sedentary lifestyle. In addition to being associated with several peripheral comorbidities including cardiovascular and metabolic disorders (Head, 2015; Malnick & Knobler, 2006; Walls et al., 2012), obesity is also associated with cognitive and neurobiological dysfunctions (Francis & Stevenson, 2013; Wang et al., 2016). Previous studies have demonstrated that obesity is associated with deficits in episodic and spatial memories (for reviews see Francis & Stevenson, 2013; Martin & Davidson, 2014; Sellbom & Gunstad, 2012; Yeomans, 2017) but also with increased emotional responses and affective disorders (Mansur et al., 2015).

The prevalence of obesity has also risen in young people (Ogden et al., 2016; Sahoo et al., 2015). Recent studies indicate indeed that obese adolescents display blunted performance in geometric/visuospatial problems or relational memory (Khan et al., 2015; Nyaradi et al., 2014; Øverby et al., 2013). Given that childhood and adolescence are crucial periods for cognitive and brain development (Spear, 2000), they represent a window of vulnerability to external insults such as the deleterious impact of various diets (for reviews see Andersen, 2003; Noble & Kanoski, 2016; Reichelt & Rank, 2017). In rodents, we recently showed that periadolescent high-fat diet (HFD), from weaning to adulthood (covering adolescence) induced complex motivational (Naneix et al., 2017; Tantot et al., 2017) and memory deficits (Boitard et al., 2012, 2014, 2015, 2016; Khazen et al., 2019). Importantly, we found that periadolescent, but not adult, HFD alters relational and spatial memories but enhances emotional memories (for reviews see Del Olmo & Ruiz-Gayo, 2018; Morin et al., 2017; Murray & Chen, 2019). However, how these memory alterations are supported by specific neurobiological changes is still unclear.

Relational and emotional memories are differently dependent on the hippocampus (Bunsey & Eichenbaum, 1996; Hartley et al., 2014) and the amygdala (LeDoux, 2003; McGaugh, 2004; Paré, 2003). In humans, clinical studies have shown that obese patients present hippocampal (Mestre et al., 2017; Mueller et al., 2012) and amygdala (Connolly et al., 2013; Pasquali et al., 2006; Widya et al., 2011) alterations. It is noticeable that both hippocampus- and amygdala-related cognitive and neurobiological processes complete their development during adolescence (for reviews see McCormick & Mathews, 2010; Saygin et al., 2015; Spear, 2000). Interestingly, overweight/obese children present reduced hippocampal volumes (Bauer et al., 2015) and increased amygdala activation (Boutelle et al., 2015). Similar patterns have been reported in HFD animal models (Abbott et al., 2019; Bose et al., 2009). Therefore, hippocampus and amygdala may then be highly vulnerable to the long-term deleterious effects of periadolescent HFD. However, the causal role of these two brain areas in periadolescent HFD-related memory changes remain to be demonstrated.

Here we investigated the causal role of ventral hippocampus (vHPC) and the basolateral amygdala (BLA) in memory deficits induced by periadolescent HFD. We previously assessed spatial and relational memory using aversive (Boitard et al., 2014, 2016) or rewarded (Boitard et al., 2012) learning tasks. Here we used different variations of non-aversive, non-rewarded, spontaneous learning tasks using objects, i.e. object recognition memory (ORM). By manipulating the delay between training and test, as well as the habituation to the training context, we used different situations that could differentially recruit the hippocampus (for review see Cohen & Stackman, 2015) and/or the BLA (Maroun & Akirav, 2008; Okuda et al., 2004; Roozendaal et al., 2006). We first showed that periadolescent HFD decreased specifically hippocampal-dependent form of long-term ORM in non-habituated rats (high arousal conditions). Using a chemogenetic DREADD approach (Armbruster et al., 2007; Rogan & Roth, 2011), we then demonstrated that this ORM deficit is abolished by the inhibition of the vHPC, but not the BLA, excitatory neurons during the acquisition. Contrastingly, we also observed that the HFD-induced enhancement of aversive odor memory is dependent on BLA, but not vHPC, activity.

## 2 METHODS

### 2.1 Animals and diets

Naïve male Wistar rats (Janvier), aged 3 weeks when they arrived, were housed in groups of two to four individuals in polycarbonate cages (48 x 26 x 21 cm) in an acclimatized (22 ± 1°C) housing room maintained under a 12 h light/dark cycle (lights on at 8:00 am, lights off at 8:00 pm). They had *ad libitum* access to food and water from their arrival until euthanasia day. At their arrival, rats were maintained either on a control diet (CD; 2.9 kcal/g; 8% lipids, 19% proteins, 73% carbohydrates; A04, SAFE) or on a high fat diet (HFD; 4.7 kcal/g; 45% lipids, 20% proteins, 35% carbohydrates; D12451, Research Diet). Animals’ body weight was recorded weekly. Rats were exposed to CD or HFD for 12 weeks (from weaning to adulthood) before the start of the behavioral experiments (**Figure 1A**). After the completion of the ORM task, rats were housed individually in identical cages (48 x 26 x 21 cm) for the duration of the conditioned odor aversion procedure.

**Figure 1.**
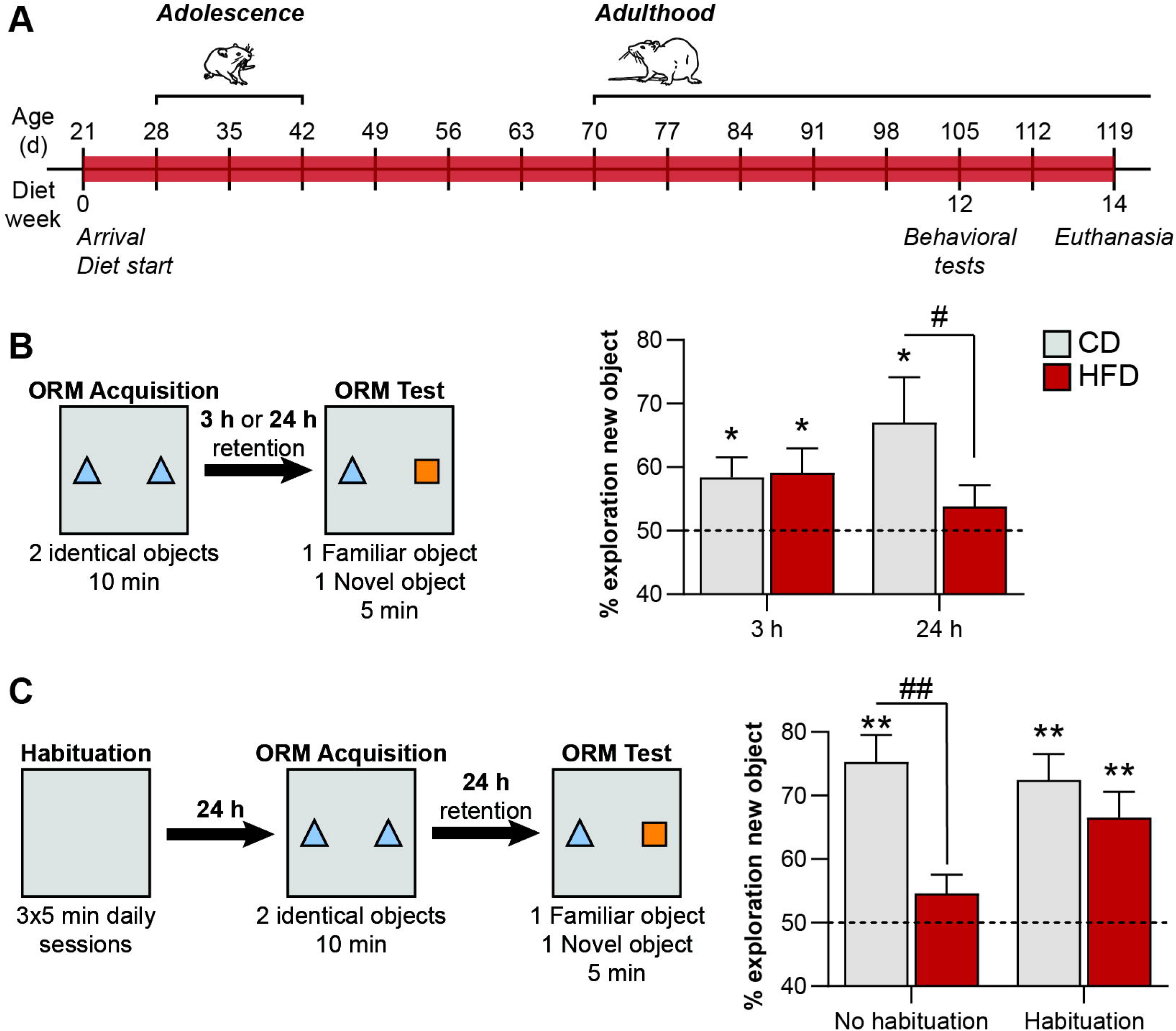
Periadolescent HFD exposure altered long-term object location memory (ORM). (A) Schematic representation of the experimental design. Rats had *ad libitum* access to either CD (grey bars) or HFD (red bars) from weaning to adulthood. All behavioral testing occurred at adulthood after at least 12 weeks of diet. (B) Periadolescent HFD exposure alters long-term (24h testing), but not short-term (3h testing) ORM. (C) Diet-induced ORM deficit at 24h is abolished when the animals were previously habituated to the arena. Data are represented as mean ± SEM. * p < 0.05, ** p < 0.01 (one sample t-test versus 50%), # p < 0.05, ## p < 0.01 (Diet effect, two-way ANOVAfollowed by Sidak’s *post hoc* tests).

All procedures were performed in agreement with the French (Directive 2013-118, 1 February 2013) and international (directive 2010-63, 22 September 2010, European Community) legislations and received approval from the local Ethics Committee (5012047-A).

### 2.2 Viral vector and drugs

An adeno-associated viral vector (AAV) carrying the inhibitory hM4D(Gi) DREADD driven by the CaMKII promoter (to limit expression to excitatory neurons) was obtained from University of North Carolina Vector core (Chapel Hill, NC, USA). The vector used was an AAV8-CaMKII-hM4D(Gi)-mCherry (3–4×10^12^ vp/ml).

The exogenous ligand Clozapine-N-Oxyde (CNO; Enzo Life Sciences) was dissolved in 0.9% saline containing 0.5% of dimethyl sulfoxide (DMSO; Sigma) at a final concentration of 1 mg/ml. Saline solution (0.9%) with 0.5% DMSO was used for vehicle injections. Both CNO and vehicle were prepared fresh for every injection day and injected (i.p.) 45 min before behavioral testing.

### 2.3 Surgery

After 7-8 weeks under CD or HFD, rats were anaesthetized under isoflurane (5% induction; 1–2 % maintenance), injected with the antalgic buprenorphine (Buprecare; 0.05 mg/kg, s.c.) and mounted on a stereotaxic apparatus (David Kopf Instruments). The scalp was shaved, cleaned and locally anaesthetized with and local application of xylocaine. The viral vector was infused using repeated pressure pulses delivered via a glass micropipette connected to a Picospritzer III (Parker, NH, USA). For the vHPC, 1 μl of the AAV was injected over 5 min (200 nl/min) at 2 sites in each hemisphere (i.e. 2 ul per hemisphere). The vHPC coordinates were AP −5.5 mm, ML ±5.5 mm from Bregma, DV −4 and −6 from the skull surface (Paxinos & Watson, 2007). For the BLA, 1 μl of the AAV was injected over 5 min at 1 site in each hemisphere: AP −3.0 mm, ML ±5.5 mm from Bregma, DV −8 mm from the skull surface. The pipette was left in place for 5 additional minutes before being slowly removed. Rats were housed in pairs immediately after surgery and were allowed at least 4 weeks to recover before the start of behavioral testing to allow ample time for virus expression.

### 2.4 Object recognition memory (ORM)

ORM is a classical procedure to assess non-spatial memory based on the recognition of a familiar object and the natural rodent’s tendency to explore novel, non-threatening, object. ORM requires a single trial and does not involve any aversive or food reward component (Ennaceur, 2010; Ennaceur & Delacour, 1988).

ORM task was performed in an arena sized 1.0 m x 1.0 m x 0.80 m (W x L X H), between 9:00 am and 1:00 pm. During the acquisition phase, rats were placed in the apparatus for 10 min and the time spent exploring two identical unfamiliar objects (either pairs of glass jars or milk cans; counterbalanced between groups) was recorded. Three or 24 h later, rats were placed back into the same apparatus containing a familiar and a novel object for 5 min at the same location than during the acquisition and the time spent exploring each object was recorded. The position of the familiar and the novel object (left or right) was counterbalanced between animals. Both objects and apparatus were cleaned with 70% of ethanol between each animal. Naïve rats usually prefer exploring the novel object, indicating memory for the familiar one, while a fail of this recall is considered as a memory deficit (Cohen & Stackman, 2015; Ennaceur, 2010; Ennaceur & Delacour, 1988). Videos were recorded for each individual rat. Object exploration was analyzed offline in blind conditions using a video tracking software (Videotrack; Viewpoint, France). Object exploration was considered when the rat was at a distance of at least 1.0-1.5 cm and moved its whiskers towards the object. Exploration values were excluded if the animal was not exploring during either the training or the testing phase, and if one object was moved during the test. Exploration is represented as the absolute time exploring each object in seconds. ORM was expressed as the percentage of exploration of the novel object during the testing phase, calculated as following: time spent exploring the novel object / (time spent exploring the novel object + time spent exploring the familiar object) x 100. A value above 50% indicates a higher exploration of the new object over the familiar one. In chemogenetic experiments, rats received either vehicle or CNO (i.p.) 45 min before the acquisition session.

In some experiments, an initial context habituation phase was performed before the acquisition session. Context habitation consisted of 3 x 5 min daily sessions during which rats were free to explore the arena without objects.

### 2.5 Conditioned odor aversion (COA)

COA results from the association of an odorized tasteless solution with a visceral malaise. In the present experiment, COA was evaluated using a previously described procedure (see Boitard et al., 2015). Rats were first acclimated to a water-deprivation regimen for 4 days. Access to water was provided in a graded bottle (with 0.5 ml accuracy) placed in the rats’ home cage for 15 min each day between 9am and 11am. Baseline water consumption was obtained by averaging the intake of the last 3 days. On the fifth day, rats had access for 15 min to almond- (0.01% benzaldehyde; Sigma Aldrich) or banana-scented (0.01% isopentyl acetate; Sigma Aldrich) water, counterbalanced between rats. The percentage of odorized solution consumption with respect to water baseline was used as a measure of neophobia. Thirty minutes after, rats received an intraperitoneal injection of lithium chloride (LiCl; Sigma Aldrich; 25 mg/kg, 0.075M, 0.75 % of body weight). On days 6 and 7, rats had access to water for 15 min each day to re-establish baseline water intake. Finally, on day 8, long-term memory of the odor aversion was assessed by providing access to the almond- or banana-odorized water for 15 min, immediately followed by 15 min of water. The percentage of odorized water consumption with respect to the initial consumption of the same solution during conditioning was used as a measure of the strength of conditioned odor aversion. In chemogenetic experiments, CD and HFD rats received either vehicle or CNO (i.p.) 45 min before the first presentation of scented water on day 5.

### 2.6 Histology

After the completion of behavioral testing, rats were deeply anaesthetized using a pentobarbital monosodic/lidocaine solution (20 mg/kg) before being transcardially perfused by ice-cold saline (0.9%) followed by 4% paraformaldehyde in 0.1 M phosphate buffer. Brains were post-fixed overnight in 4% paraformaldehyde and then switched in 0.1 M phosphate buffer saline (PBS) solution and stocked at −4 °C before slicing. Serial coronal sections (40 μm) were cut using a vibratome (VT1200S, Leica Microsystems). Free-floating sections were prepared by rinsing in 0.1 M PBS for 20 min (4 x 5 min rinses), blocked for 1 h (PBS 0.1 M, 0.2% Triton-X, 4% normal goat serum) and placed in 1:1000 rabbit anti-RFP (red fluorescent protein; PM005, CliniSciences) at 4° C for 48 h. Sections were then washed in PBS for 20 min (4 x 5 min rinses), incubated in 1:200 AffiniPure rhodamine goat anti-rabbit (11-025-003; Jackson Immunoresearch) diluted in PBS for 2 h at room temperature and counterstained with 1:5000 Hoestch solution (bisBenzimide H 33258, Sigma-Aldrich). Sections were washed for 20 min in PBS (4 x 5 min rinses), mounted, and cover-slipped with Fluoromount-G (SouthernBiotech). Sections were imaged using a Nanozoomer slide scanner and analyzed with the NDP.view 2 freeware (Hamamatsu Photonics, Bordeaux Imaging Center).

### 2.7 Data analysis

Weight was analyzed using two-way repeated measures ANOVA using Diet (CD or HFD) as between-subject factor and Week of diet as within-subject factor, followed by Bonferroni’s *post hoc* tests. Total exploration time and the percentage of exploration of the novel object were analyzed using either one-way ANOVA using Group as between-subject factor or two-way repeated measures ANOVA using Diet (CD, HFD) as between-subject factor and Retention time (3h, 24h) or Habituation (with, without) as within-subject factor, followed by Sidak’s *post hoc* tests. In case of missing values in the behavioral results from the ORM task (e.g. animal excluded for an absence of exploratory behavior), a Mixed-effect model was used instead for repeated measures analyses. For chemogenetic experiments, ORM and COA measures were analyzed using one-way ANOVA with Group as between-subject factor, followed by Dunnet’s *post hoc* test versus CD-Vehicle group. Planned comparisons analyses restricted to the HFD groups (Sidak’s corrected multiple t-tests) were used to analyzed specifically chemogenetic effects. Statistical analyses were carried out on GraphPad Prism version 7. All values were expressed as mean ± standard error of the mean (SEM). The alpha risk of rejection of the null hypothesis was 0.05.

## 3 RESULTS

### 3.1 HFD intake significantly induces overweight

Upon arrival, animals were randomly divided to create two groups of similar body weight, and then exposed to either CD or HFD. After 6 and 12 weeks, rats fed with HFD were significantly heavier than CD animals (2-way repeated measures ANOVA: Diet x Diet F_2, 154_ = 31.1, p<0.001; Sidak’s *post hoc* test: Week 0, p = 0.99; Week 6 and 12 all p < 0.001; **Table 1**) as previously reported (Boitard et al., 2012, 2014, 2015, 2016).

**Table 1:** Effects of Control (CD; n = 38) and periadolescent High Fat (HFD, n = 41) diet on rats’ body weight (g). Diet started at weaning (3-weeks old, Week 0) and lasted at least 12 weeks before the start of behavioral testing. *** p < 0.001 Diet effect (two-way ANOVA followed by Bonferroni’s *post hoc* tests).

| Week | CD Body weight (g) | HFD Body weight (g) |
| --- | --- | --- |
| 0 | 74.1 ± 1.0 | 76.4 ± 1.0 |
| 6 | 392.0 ± 5.1 | 431.4 ± 5.4*** |
| 12 | 495.9 ± 7.5 | 575.4 ± 9.5*** |

### 3.2 HFD intake induces long-term ORM deficits when training takes place in a novel context

Without previous habituation to the arena, CD (n = 11) and HFD-fed (n = 17) rats were placed into it with two identical objects (**Figure 1B**). During the acquisition phase, all groups similarly explored the two identical objects (**Supplemental Figure 1A**; Diet F_1,26_ = 1.4, p =0.2; Diet x Retention Time F_1,18_ = 5.9, p = 0.03). Three or 24 h after training, rats were exposed to a familiar and a novel object. When tested 3 h after the acquisition phase, CD- and HFD-fed rats exhibited a similar higher exploration of the novel object (**Figure 1B**; one-sample t-test versus 50%: CD t_9_ = 2.7, p = 0.026; HFD t_16_ = 2.4, p = 0.03). However, when tested 24 h after the acquisition, only CD rats showed a significant preference for the novel object (one-sample t-test versus 50%: CD t_7_ = 2.4, p = 0.047; HFD t_13_ = 1.2, p = 0.3). Mixed-effect analysis confirmed a significant Diet x Retention time interaction (F_1,18_ = 4.9, p = 0.04; Diet F_1,26_ = 3.9, p = 0.06). More important, *post hoc* analyses confirmed a significant difference between the two diet groups when tested at 24 h (p < 0.05) but not at 3 h (p > 0.9; Sidak’s *post hoc* tests), indicating that HFD consumption during adolescence impairs long-term, but not short-term, ORM.

We then evaluated whether previous habituation to the context would alleviate HFD-induced long-term ORM deficits. For this purpose, another batch of CD (n = 6) and HFD-fed (n = 7) rats was tested twice in the ORM task at 24h after training, first without exposure to the training arena to confirm previous diet effect and then with habituation to a novel training context (**Figure 1C**). In both conditions, the two diet groups similarly explored the two objects during the acquisition phase (**Supplemental Figure 1B**; Diet t_11_ = 0.04, p = 1.0). However, the exposure to the arena increased the time exploring the novel object selectively in the HFD group (two-way repeated measures ANOVA: Diet F_1,11_ = 8.8, p = 0.01; Diet x Habituation F_1,11_ = 5.8, p =0.04). As previously observed, HFD rats tested without being habituated to the training arena presented a lower exploration of the new object compared to the CD group (p < 0.01; Sidak’s *post hoc* test), replicating a deficit in long-term ORM (one-sample t-test versus 50%: CD, t_5_ = 6.0, p = 0.001; HFD, t_6_ = 1.6, p = 0.2). However, after having been exposed to a new arena for three consecutive days before the acquisition phase, the same animals significantly preferred to explore the novel object than the familiar one (one-sample t-test versus 50%: CD, t_5_ = 5.5, p = 0.003; HFD, t_5_ = 4.1, p = 0.007), similarly to the CD rats (p = 0.6), demonstrating in this case an intact long-term ORM.

### 3.3 Chemogenetic inactivation of the ventral hippocampus, but not the basolateral amygdala, rescues HFD-induced recognition memory impairment

We then investigated the contribution of the vHPC and the BLA in periadolescent-HFD induced memory alterations, using a chemogenetic approach involving the targeted expression of the inhibitory DREADD hM4Di in excitatory neurons thanks to the CaMKII promoter (**Figure 2**). After histological analyses, HFD rats presenting bilateral hM4Di-mCherry expression in either the vHPC (n = 15) or in the BLA (n = 13) were kept for the statistical analyses. These rats were further divided depending if they received vehicle (vHPC, n = 8; BLA, n = 8) or CNO injection (HFD-vHPC-CNO, n = 7; HFD-BLA-CNO, n = 5). An additional group of HFD-fed rats did not receive any virus injection to control for the specificity of the CNO effects on behavioral measures (n=26). They were either injected with vehicle (n=11) or CNO (HFD-No DREADD-CNO, n = 15). The HFD-fed groups injected with vehicle (with DREADD, n = 16; without DREADD, n= 11) were pooled to form a HFD group receiving vehicle (HFD-Vehicle, n = 27). Finally, control CD (with or without DREADD) rats which received vehicle injections were pooled to provide a single CD-Vehicle group (n = 21). To summarize, the statistical analyses were performed on the following 5 final groups: CD-Vehicle, n = 21; HFD-Vehicle, n = 27; HFD-No DREADD-CNO, n = 15; HFD-vHPC-CNO, n = 7; HFD-BLA-CNO, n = 5.

**Figure 2.**
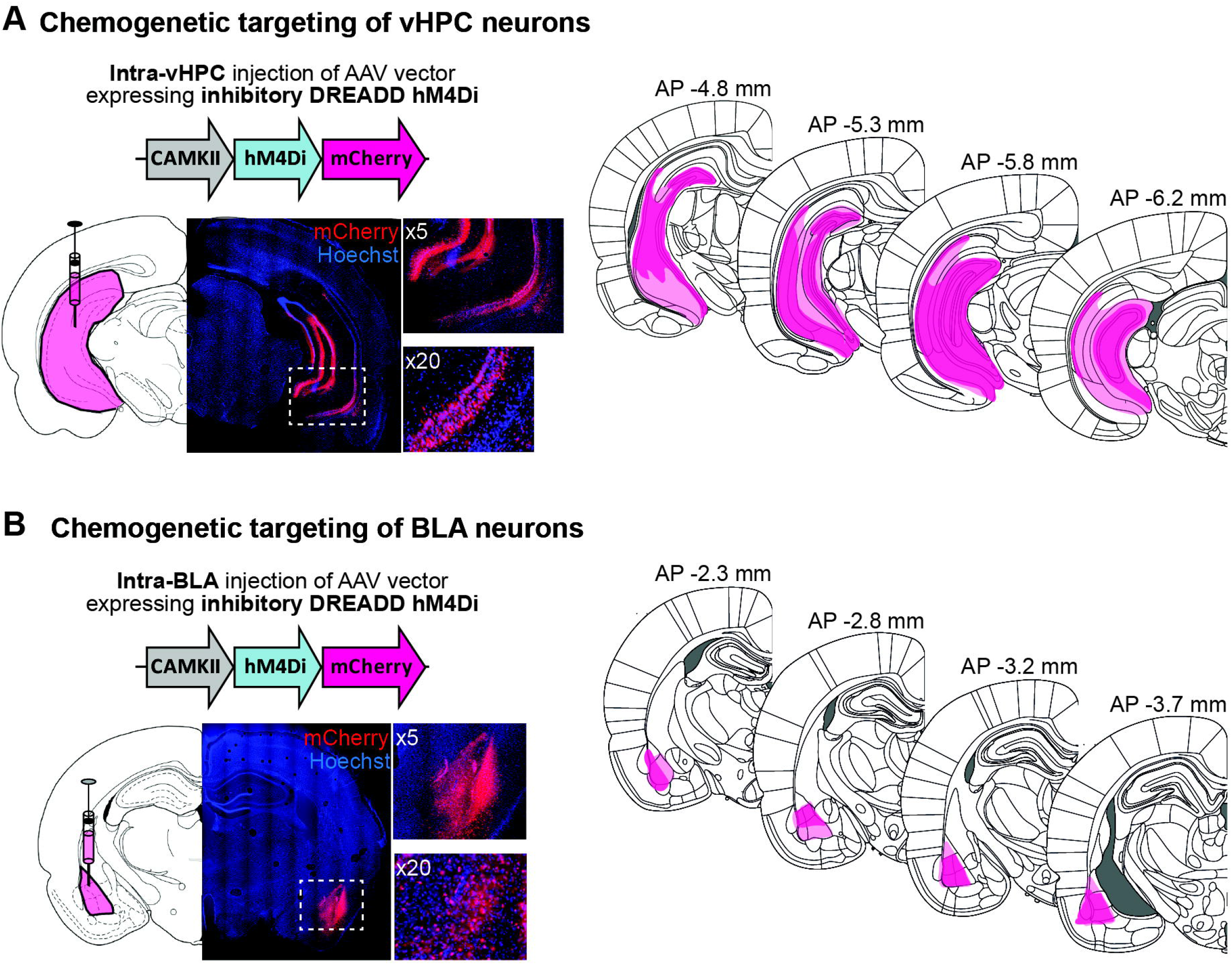
Chemogenetic targeting of the vHPC (A) or the BLA (B). *Left* Representative images illustrating the placement of AAV8-CaMKII-hM4Di-mCherry expression in the vHPC. Insets represent magnification of the area of interest (white square). *Right* Schematics adapted from Paxinos and Watson (2013) showing the largest (light pink) and smallest (dark pink) viral infection for rats included in ORM and COAexperiments.

Four weeks after surgery, long-term ORM was tested without habituation using the exact same procedure as previously described to demonstrate HFD-induced deficits (**Figure 3A**). All rats received an injection of either CNO or its vehicle 45 min before the acquisition phase. CNO injection did not alter the exploration of the objects during this phase (**Supplemental Figure 1C;** F_4,70_ = 2.2, p = 0.08). Whereas HFD-fed rats treated with Vehicle exhibited an absence of long-term ORM when tested 24 h after the acquisition, all HFD groups which received CNO showed a higher exploration of the novel object (**Figure 3B**; one-sample t-test versus 50%; CD-Vehicle t_20_ = 7.8, p < 0.001; HFD-Vehicle t_26_ = 1.9, p = 0.07; HFD-No DREADD-CNO t_14_ = 4.6, p < 0.001; HFD-vHPC-CNO t_5_ = 6.7, p < 0.001; HFD-BLA-CNO t_4_ = 4.1, p = 0.02). One-way ANOVA indicated a significant difference between the five groups (F_4,70_ = 9.0, p < 0.001). *Post hoc* analyses confirmed the significant difference in ORM performance between the CD control group and HFD-Vehicle treated rats (p < 0001) but with the other HFD groups (all p > 0.3; Dunnett’s *post hoc* test versus CD-Vehicle). More importantly, planned comparisons restricted the HFD groups confirmed the higher exploration of the novel object in the HFD-vHPC-CNO group (p < 0.001; Sidak’s multiple planned comparisons) demonstrating the reversal of ORM deficits by the vHPC inhibition during the acquisition phase. These analyses also showed an effect of CNO on memory performance in HFD-No DREADD-CNO compared to vehicle-treated HFD rats (p = 0.04), suggesting a potential unspecific CNO effect, which remained smaller than the effect observed in HFD-vHPC rats (HFD-No DREADD-CNO versus HFD-vHPC-CNO, p = 0.05). Moreover, this CNO effect was not observed in HFD-BLA group (p = 0.7 versus HFD-Vehicle; p = 0.9 versus HFD-No DREADD-CNO).

**Figure 3.**
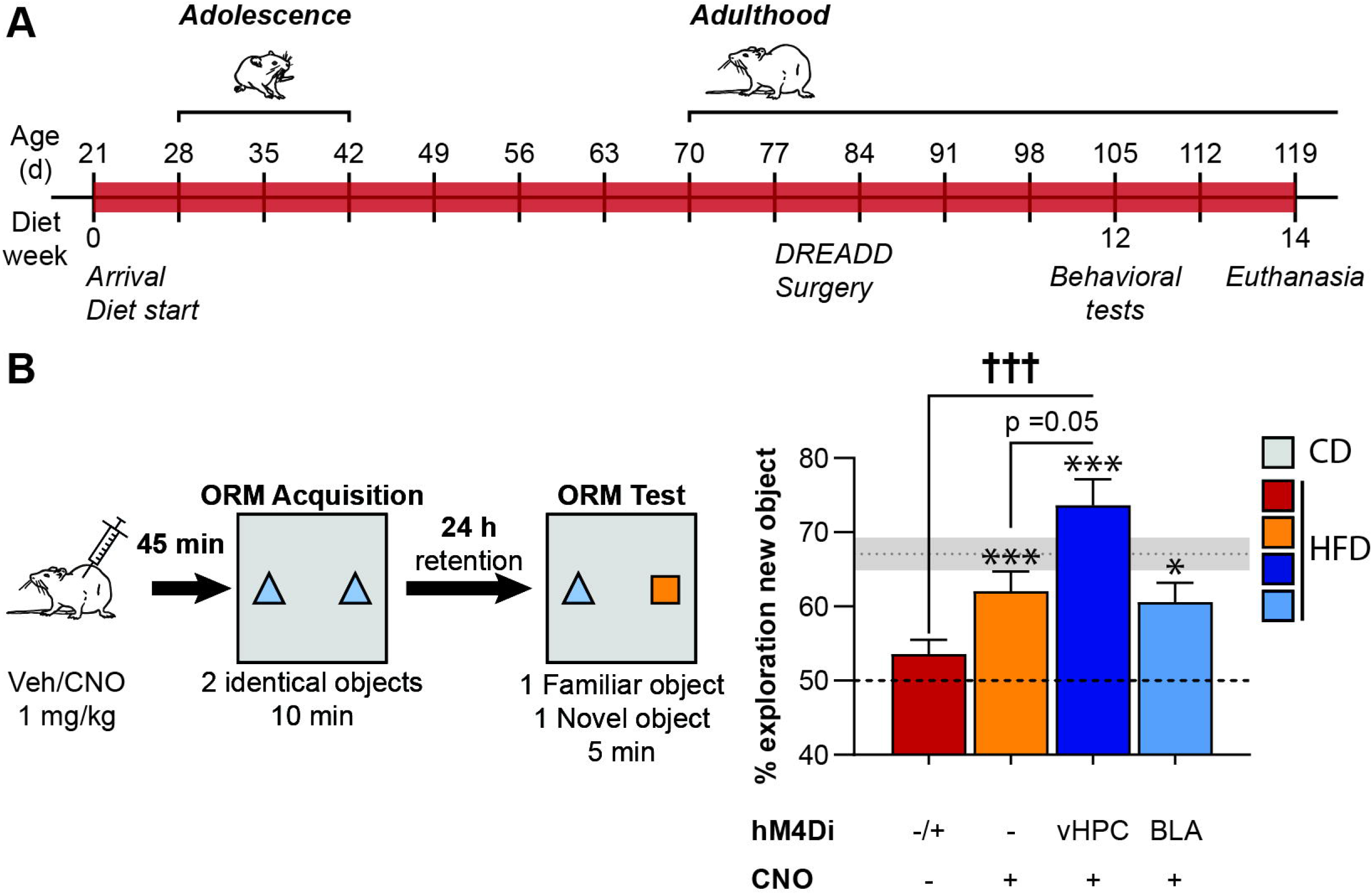
Chemogenetic inhibition of the vHPC, but not the BLA, restored long-term ORM induced by periadolescent HFD exposure. (A) Schematic representation of the experimental design. Rats had *ad libitum* access to either CD or HFD from weaning to adulthood. DREADD surgery was performed at adulthood (7-8 weeks of diet) and rats recovered 4 weeks before the start of behavioral testing. (B) Long-term ORM performance in HFD-fed rats treated with vehicle (red bars; ± indicating with or without DREADD expression), CNO but without DREADD expression (orange bars), or CNO with DREADD expressed in the vHPC (dark blue) or the BLA (light blue) Expression of the inhibitory DREADD hM4Di is depicted by structure (vHPC or BLA), except for the vehicle group. Data are represented as mean ± SEM. Shaded grey zone represents ORM performance of the CD-Vehicle group ± SEM. * p< 0.05, ** p < 0.01 (one sample t-test versus 50%), ††† p < 0.001 (Group effect, Sidak’s multiple comparisons test).

### 3.4 Chemogenetic inactivation of the basolateral amygdala, but not the ventral hippocampus, rescues HFD-induced aversion memory enhancement

We investigated the impact of chemogenetic inhibition of vHPC and BLA excitatory neurons on the enhancement of aversion memory induced by periadolescent HFD exposure using a COA procedure (**Figure 4A**; CD-Vehicle, n = 18; HFD-Vehicle, n = 23; HFD-No DREADD-CNO, n = 9; HFD-vHPC-CNO, n = 7; HFD-BLA, n = 5). Neither the diet nor the injection of CNO/vehicle affected the consumption of odorized water during its first presentation, i.e. odor-malaise association (**Figure 4B**; one-way ANOVA Group F_4,57_ = 0.9, p = 0.5), and all groups presented a low level of neophobia toward the new odorized water (all p > 0.09; one-sample t-test versus 100% water baseline). However, the long-term aversion memory was differently impacted by periadolescent HFD and chemogenetic inhibition of the vHPC or the BLA during the odor-malaise pairing (**Figure 4C**; one-way ANOVA Group F_4,57_ = 3.9, p = 0.007). As previously observed (Boitard et al., 2015, 2016), periadolescent HFD exposure induced a stronger aversion memory compared to the control CD group (p = 0.05, Dunnett’s *post hoc* test). However, the relative consumption of odorized water of CNO-treated HFD groups is similar to the CD one (all p > 0.2, Dunnett’s *post hoc* test). Restricted analysis to the HFD groups showed that neither the injection of CNO alone nor vHPC inhibition during odor-malaise pairing reduced the aversion memory (all p > 0.9, Sidak’s multiple planned comparisons test compared to HFD-Vehicle). However, the previous inhibition of the BLA significantly reduces the aversion for the odorized solution (p < 0.01), demonstrating that BLA activity during the conditioning is necessary for periadolescent HFD-induced aversive memory enhancement.

**Figure 4.**
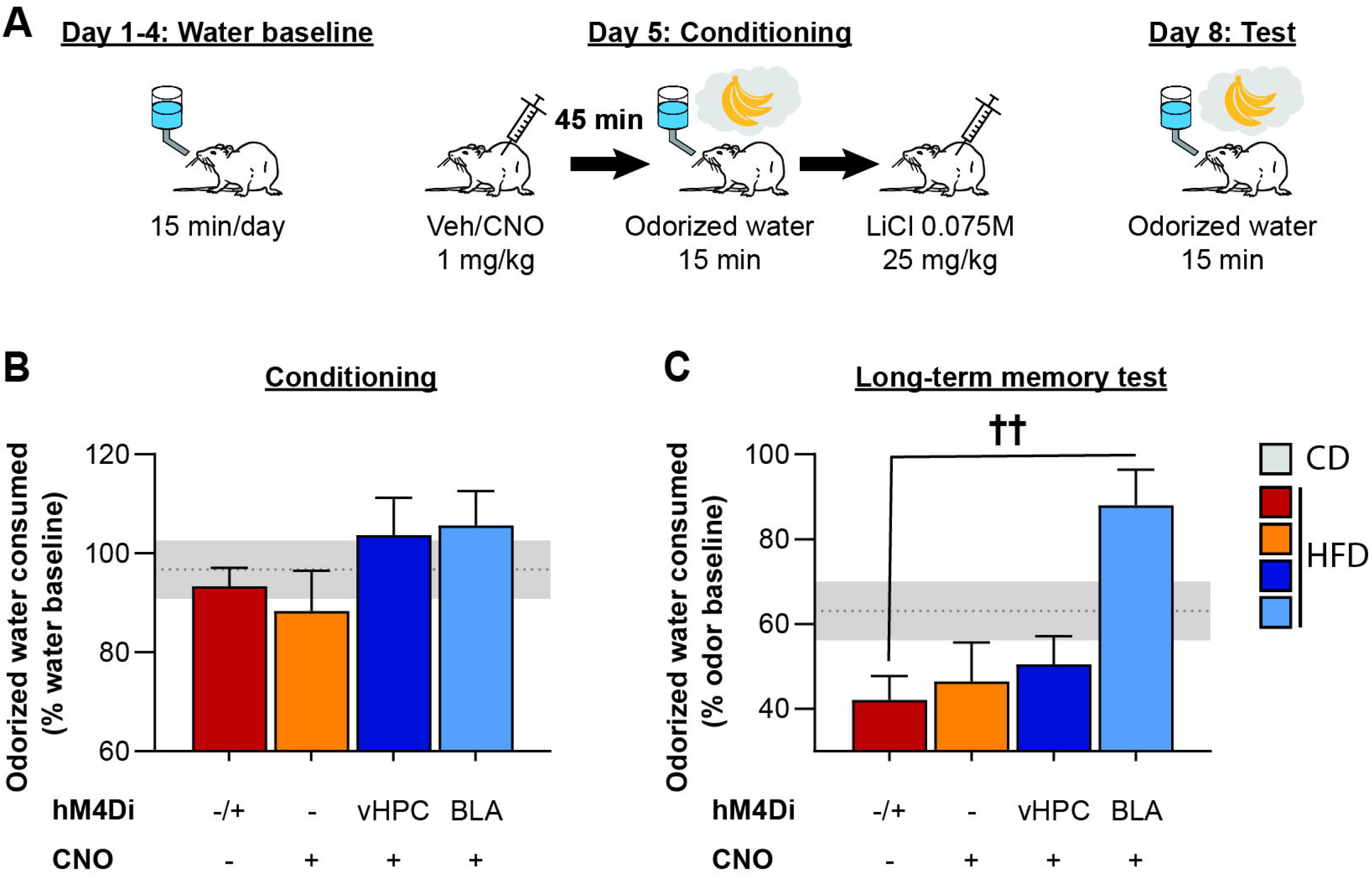
Chemogenetic inhibition of the BLA, but not the vHPC, rescued enhanced aversion memory induced by periadolescent HFD exposure. (A) Schematic representation of conditioned odor aversion (COA) protocol and chemogenetic inhibition of vHPC or BLA before COA conditioning. (B) Neither HFD or CNO injection impacted the consumption of odorized water during the conditioning phase (in percentage of water baseline consumption). (C) Long-term COA memory in HFD-fed rats treated with vehicle (red bars; ± indicating with or without DREADD expression), CNO but without DREADD expression (orange bars), or CNO with DREADD expressed in the vHPC (dark blue) or the BLA (light blue). Expression of the inhibitory DREADD hM4Di is depicted by structure (vHPC or BLA), except for the vehicle group. Data are represented as mean ± SEM. Shaded grey zone represents ORM performance of the CD-Vehicle group ± SEM. †† (Group effect, Sidak’s multiple comparisons test).

## 4 DISCUSSION

Here, we demonstrated that periadolescent HFD consumption (from weaning to adulthood) induced long term memory alterations. Specifically, we showed that HFD-fed rats presented a deficit in long-term, but not short-term, ORM when they are exposed to a novel context. Using chemogenetic inactivation, we found that manipulation of the vHPC, but not of the BLA, restored HFD-induced long-term ORM deficit. On the contrary, chemogenetic silencing of the BLA, but not of the vHPC, normalized the increased aversive memory observed in HFD-fed animals.

### 4.1 Effects of periadolescent high-fat diet on object recognition memory

The effects of HFD on object-related memory has led to contradictory results (for reviews see Abbott et al., 2019; Cordner & Tamashiro, 2015). The present study indicates that 12 weeks of exposure to HFD, starting at weaning, is sufficient to alter rats’ object memory but only under certain conditions. Indeed, we observed that periadolescent HFD impaired long-term ORM tested 24h after sampling novel objects in a novel context, but had no effect on short-term ORM tested 3 hours after training. These results are consistent with previous studies indicating no effect of HFD on short-term ORM (Beilharz et al., 2014, 2016; Kendig et al., 2019; Kosari et al., 2012; Lavin et al., 2011; McLean et al., 2018; Tran & Westbrook, 2015, 2017, 2018; Tucker et al., 2012), but an impairment of long-term object memory (Ayabe et al., 2018; de Andrade et al., 2017; Mucellini et al., 2019; Wang et al., 2016; Zuloaga et al., 2016). This differential impact of diet suggests a specific effect of peridadolescent HFD on memory consolidation processes. Interestingly, we previously reported a similar effect on consolidation of spatial memories (Boitard et al., 2014, 2016) and emotional memories (Boitard et al., 2015), but also of object location memory (Khazen et al., 2019), which suggests that HFD consumption during early life periods could interfere with cellular substrates specifically involved in memory consolidation.

Importantly, we also identified that habituation to the arena for three days prevented HFD-induced long-term ORM deficits. This could explain the absence of HFD-induced long-term ORM deficit reported in the literature after habituation and/or repeated tests (Heyward et al., 2012, 2016; Tucker et al., 2012). Prior habituation and exploration of the arena is known to reduce the processing of contextual information during memory consolidation (Cohen & Stackman, 2015; Oliveira et al., 2010) and the arousal component of the task (Maroun & Akirav, 2008; Okuda et al., 2004; Roozendaal et al., 2006). Then, the specific diet-induced deficit of long-term ORM reported here may be supported by differential neurobiological substrates involved in multiple memory systems, particularly the hippocampus and the amygdala.

### 4.2 Effects of chemogenetic manipulation of vHPC and BLA on periadolescent HFD-induced object recognition memory deficits

The hippocampus and the amygdala play a crucial role in long-term ORM (Cohen & Stackman, 2015; Roozendaal et al., 2008) and are profoundly affected by exposure to HFD during the periadolescent period (Del Olmo & Ruiz-Gayo, 2018; Morin et al., 2017; Murray & Chen, 2019; Reichelt, 2016; Reichelt & Rank, 2017). We therefore wondered whether chemogenetic inhibition of these two brain regions could alleviate the diet effects on object memory.

Recent studies have suggested that CNO may be metabolized *in vivo* to clozapine, an atypical antipsychotic drug, able to interact with DREADD receptors but also to induce non-DREADD related effects (Gomez et al., 2017; Ilg et al., 2018; MacLaren et al., 2016). To rule out this possibility, we included a control group which received CNO at a dose known to induce marginal behavioral effect. Our results showed a slight improvement in long-term ORM in this group which suggests that metabolism of CNO to clozapine might induce some behavioral effects as shown following clozapine administration (Addy et al., 2005; Mutlu et al., 2011). This pattern of results in the control group cannot however account for the complete restoration of ORM deficits following chemogenetic inhibition of vHPC, hence highlighting the central role of this brain region in HFD-induced long-term ORM deficits. Such results are in agreement with previous research which has demonstrated that long-term ORM, but not short term ORM, relies on the hippocampus (for review see Cohen & Stackman, 2015). Moreover, previous studies have shown that hippocampal manipulations have a greater impact on long-term ORM when performed in an unfamiliar context (Kim et al., 2014; Oliveira et al., 2010), whereas the perirhinal cortex is crucial in both familiar and unfamiliar contexts (Kim et al., 2014). These results suggest that, in a novel context, novel objects may be encoded as part of the context thereby involving the hippocampus, whereas if the novel objects are presented in a familiar environment they are encoded under a process that probably does not involve contextual information processing and therefore does not rely on the hippocampus. We could then hypothesize that HFD-fed animals did not exhibit a memory deficit when they were previously habituated to the arena, as in that case ORM performance is not dependent on a dysfunctional hippocampus.

Habituation to the training context also greatly influences the impact of emotional arousal and the BLA in long-term ORM (Maroun & Akirav, 2008; Roozendaal et al., 2006). However, chemogenetic BLA silencing did not have a greater effect than those of CNO alone on HFD-induced ORM deficit. The amygdala is one of the major target of vHPC projection neurons (Pitkänen et al., 2000), suggesting that the vHPC-to-BLA pathway is not involved in the beneficial effect of silencing vHPC excitatory neurons on long-term ORM. The vHPC involvement in memory processes also involves other projections to the nucleus accumbens or the ventromedial prefrontal cortex (Barker et al., 2019; Hsu et al., 2018; Okuyama et al., 2016; Phillips et al., 2019) and future studies are warranted to determine the role of these circuits in HFD-induced memory deficits.

### 4.3 Effects of chemogenetic manipulation of BLA and vHPC on periadolescent HFD-induced aversive memory enhancement

Few studies have examined the effects of HFD on aversive memory. Aversive cue-based memory is highly dependent on the BLA (LeDoux, 2003; McGaugh, 2004; Paré, 2003). We previously found that periadolescent HFD enhanced long-term, but not short-term, odor aversion memory as well as long-term auditory fear memory (Boitard et al., 2015, 2016). Here we replicate this finding and we provide evidence that chemogenetic silencing of the BLA, but not the vHPC, normalized the increased odor aversion memory observed in the HFD group. It is noticeable that, contrary to the ORM, CNO injection alone (without any DREADDs) did not have any effect by itself on HFD-induced aversive memory enhancement.

It is generally considered that during emotional arousal the activity of the BLA is modulated by glucocorticoids and noradrenaline, and eventually impacts aversive memory via glutamatergic projections to other structures, including the hippocampus (McEwen et al., 2016; McGaugh, 2004). In this context, we previously found that blockade of glucocorticoid receptors in the BLA is able to normalise the enhanced aversive memory of HFD group (Boitard et al., 2015). Taken together, these data suggest that periadolescent HFD consumption increases the activation of BLA excitatory neurons through glucocorticoids in response to emotional experience, leading to an enhanced odor aversion memory. Furthermore, a recent study showed that chemogenetic inactivation of the noradrenergic pathway from the locus coeruleus to the BLA abolished aversive memory enhancement, but not ORM impairment, induced by chronic pain (Llorca-Torralba et al., 2019). According to the differential effect of chemogenetic BLA silencing on aversive memory and ORM in HFD-fed rats, a similar impact of periadolescent HFD on the noradrenergic modulation of BLA may also be involved.

In contrast, chemogenetic inactivation of the vHPC did not modify odor aversion memory in HFD-fed animals. Even though the BLA is highly connected to the vHPC (Pitkänen et al., 2000) and that the BLA-to-vHPC pathway plays a central role in emotional processes (Beyeler et al., 2016; Felix-Ortiz et al., 2013; Rei et al., 2015), our results suggest that the HFD-induced enhancement of aversive memories may rather involve projections to the nucleus accumbens (Beyeler et al., 2016; Stuber et al., 2011) or to the ventromedial prefrontal cortex (Burgos-Robles et al., 2017; Felix-Ortiz et al., 2013).

Altogether our results demonstrate in periadolescent HFD-fed rats, that silencing vHPC, but not BLA, improves long-term ORM deficits, while silencing BLA, but not vHPC, normalizes COA enhancement. This double dissociation suggests that vHPC and BLA, though related structures, can have distinct and independently-driven functions in HFD-induced memory changes.

### 4.4 Conclusions

The adolescent brain is highly sensitive and prone to cognitive alterations promoted by diets rich in fat and/or sugar (for reviews see Del Olmo & Ruiz-Gayo, 2018; Morin et al., 2017; Murray & Chen, 2019; Noble & Kanoski, 2016; Reichelt, 2016; Reichelt & Rank, 2017). Our study demonstrates that periadolescent HFD alters long-term memory processes, impairing recognition memory through vHPC-dependent processes while enhancing emotional memory through BLA specific effects. Such bidirectional effect on hippocampal and amygdala memory functions have also been reported in chronic stress and post-traumatic stress disorder (Elzinga & Bremner, 2002; Kaouane et al., 2012; Layton & Krikorian, 2002; Mahan & Ressler, 2012). Interestingly, obesity is linked to a higher prevalence of post-traumatic stress disorder, especially during adolescence (Pagoto et al., 2012; Perkonigg et al., 2009). Future investigation is necessary to evaluate how HFD consumption during adolescence impacts preferentially the medial temporal lobe, and how the potential alterations of specific hippocampal and amygdala circuits may mediate the cognitive impact of juvenile obesity.

## 5 AUTHORS CONTRIBUTION

C.B.B, G.P.L., E.C. and G.F. acquired funding; F.N., E.C. and G.F. designed research; F.N., I.B., M.S.Z. performed research; F.N., I.B., E.C. and G.F. analysed data; E.C. and G.F. supervised research; F.N., I.B., E.C. and G.F. wrote the manuscript; C.B.B, M.S.Z. and G.P.L. edited and approved the manuscript.

## 6 ACKNOWLEDGMENTS

The microscopy was completed at the Bordeaux Imaging Center, a service unit of CNRS-INSERM and Bordeaux University and member of the national infrastructure, France BioImaging. We thank María-José Olvera for technical assistance and Yoan Salafranque for the care provided to the animals during experiments.

## 8 ANNEXE 1

OBETEEN Consortium: Guillaume Ferreira^1^,*, Gustavo Pacheco-Lopez^2^,**, Etienne Coutureau^3^, Ranier Gutierrez^4^, Pascal Barat^1,5^, Federico Bermudez-Rattoni^6^, Gwenaelle Catheline^3^, Claudia I. Pérez^4^, Pauline Lafenêtre^1^, Daniel Osorio-Gomez^6^, Kioko Guzman-Ramos^2^, Fabien Naneix^1,3^, Ernesto Sanz-Arigita^1,3^, Ioannis Bakoyiannis^1^

^1^Univ. Bordeaux, INRAE, Bordeaux INP, Nutrition and Integrative Neurobiology, UMR 1286, Bordeaux, France; ^2^Metropolitan Autonomous University (UAM), Campus Lerma, Health Sciences Department, Lerma, Mexico; ^3^Univ. Bordeaux, CNRS, INCIA, UMR 5287, Bordeaux, France; ^4^Center for Research and Advanced Studies (CINVESTAV), Department of Pharmacology, Mexico City, Mexico; ^5^CHU Bordeaux, Children hospital, Bordeaux, France; ^6^National Autonomous University of Mexico (UNAM), Cellular Physiology Institute, Mexico City, Mexico. *Responsible for the consortium at the French National Research Agency (ANR). **Responsible for the consortium at the Mexican National Council for Science and Technology (CONACYT).

Note: Consortium author list recognizes the significant and irreducible commitment with the conceptualization, management, and realization of this research project.

---

**Supplemental Figure 1.**
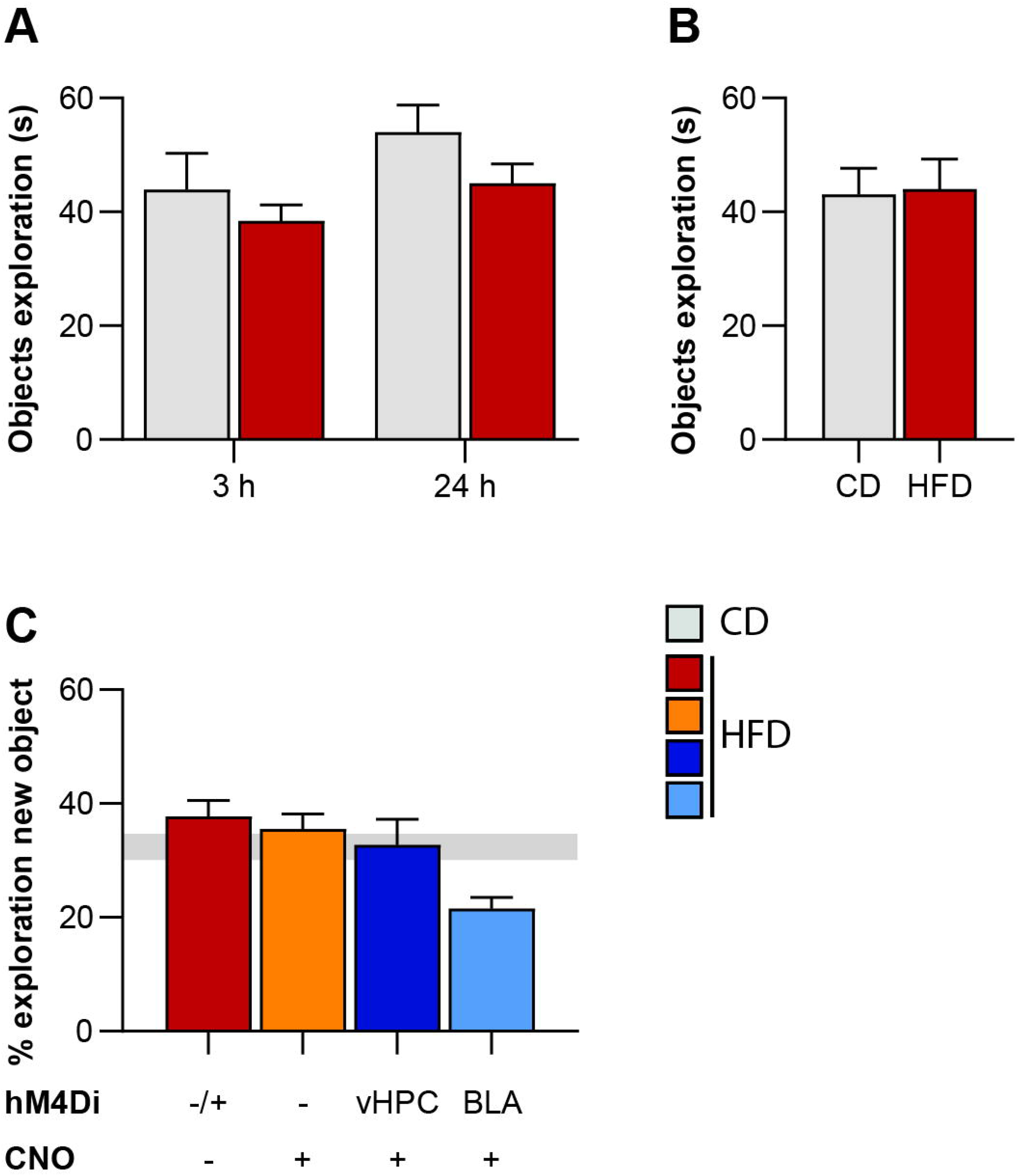
Periadolescent HFD did not alter object exploration during ORM acquisition. Total object exploration during the acquisition phase of the ORM task for CD and HFD groups being tested at 3 or 24 h without context habituation (A), being tested 24 h with context habituation (B) or receiving CNO/Vehicle injection prior to acquisition (C). Expression of the inhibitory DREADD hM4Di is depicted by structure (vHPC or BLA), except for the vehicle group (± indicating with or without DREADD expression). Data are represented as mean ± SEM.

## REFERENCES

Abbott, K. N., Arnott, C. K., Westbrook, R. F., & Tran, D. M. D. (2019). The effect of high fat, high sugar, and combined high fat-high sugar diets on spatial learning and memory in rodents: A meta-analysis. Neuroscience and Biobehavioral Reviews, 107, 399–421. https://doi.org/10.1016/j.neubiorev.2019.08.010

Addy, N. A., Pocivavsek, A., & Levin, E. D. (2005). Reversal of clozapine effects on working memory in rats with fimbria-fornix lesions. Neuropsychopharmacology: Official Publication of the American College of Neuropsychopharmacology, 30(6), 1121–1127. https://doi.org/10.1038/sj.npp.1300669

Andersen, S. L. (2003). Trajectories of brain development: point of vulnerability or window of opportunity? Neuroscience and Biobehavioral Reviews, 27(1–2), 3–18. https://doi.org/10.1016/s0149-7634(03)00005-8

Armbruster, B. N., Li, X., Pausch, M. H., Herlitze, S., & Roth, B. L. (2007). Evolving the lock to fit the key to create a family of G protein-coupled receptors potently activated by an inert ligand. Proceedings of the National Academy of Sciences of the United States of America, 104(12), 5163–5168. https://doi.org/10.1073/pnas.0700293104

Ayabe, T., Ohya, R., Kondo, K., & Ano, Y. (2018). Iso-α-acids, bitter components of beer, prevent obesity-induced cognitive decline. Scientific Reports, 8(1), 4760. https://doi.org/10.1038/s41598-018-23213-9

Barker, J. M., Bryant, K. G., & Chandler, L. J. (2019). Inactivation of ventral hippocampus projections promotes sensitivity to changes in contingency. Learning & Memory (Cold Spring Harbor, N.Y.), 26(1), 1–8. https://doi.org/10.1101/lm.048025.118

Bauer, C. C. C., Moreno, B., González-Santos, L., Concha, L., Barquera, S., & Barrios, F. A. (2015). Child overweight and obesity are associated with reduced executive cognitive performance and brain alterations: a magnetic resonance imaging study in Mexican children. Pediatric Obesity, 10(3), 196–204. https://doi.org/10.1111/ijpo.241

Beilharz, J. E., Maniam, J., & Morris, M. J. (2014). Short exposure to a diet rich in both fat and sugar or sugar alone impairs place, but not object recognition memory in rats. Brain, Behavior, and Immunity, 37, 134–141. https://doi.org/10.1016/j.bbi.2013.11.016

Beilharz, J. E., Maniam, J., & Morris, M. J. (2016). Short-term exposure to a diet high in fat and sugar, or liquid sugar, selectively impairs hippocampal-dependent memory, with differential impacts on inflammation. Behavioural Brain Research, 306, 1–7. https://doi.org/10.1016/j.bbr.2016.03.018

Beyeler, A., Namburi, P., Glober, G. F., Simonnet, C., Calhoon, G. G., Conyers, G. F., Luck, R., Wildes, C. P., & Tye, K. M. (2016). Divergent Routing of Positive and Negative Information from the Amygdala during Memory Retrieval. Neuron, 90(2), 348–361. https://doi.org/10.1016/j.neuron.2016.03.004

Boitard, C., Cavaroc, A., Sauvant, J., Aubert, A., Castanon, N., Layé, S., & Ferreira, G. (2014). Impairment of hippocampal-dependent memory induced by juvenile high-fat diet intake is associated with enhanced hippocampal inflammation in rats. Brain, Behavior, and Immunity, 40, 9–17. https://doi.org/10.1016/j.bbi.2014.03.005

Boitard, C., Etchamendy, N., Sauvant, J., Aubert, A., Tronel, S., Marighetto, A., Layé, S., & Ferreira, G. (2012). Juvenile, but not adult exposure to high-fat diet impairs relational memory and hippocampal neurogenesis in mice. Hippocampus, 22(11), 2095–2100. https://doi.org/10.1002/hipo.22032

Boitard, C., Maroun, M., Tantot, F., Cavaroc, A., Sauvant, J., Marchand, A. R., Layé, S., Capuron, L., Darnaudery, M., Castanon, N., Coutureau, E., Vouimba, R. M., & Ferreira, G. (2015). Juvenile obesity enhances emotional memory and amygdala plasticity through glucocorticoids. The Journal of Neuroscience: The Official Journal of the Society for Neuroscience, 35(9), 4092–4103. https://doi.org/10.1523/JNEUROSCI.3122-14.2015

Boitard, C., Parkes, S. L., Cavaroc, A., Tantot, F., Castanon, N., Layé, S., Tronel, S., Pacheco-Lopez, G., Coutureau, E., & Ferreira, G. (2016). Switching Adolescent High-Fat Diet to Adult Control Diet Restores Neurocognitive Alterations. Frontiers in Behavioral Neuroscience, 10, 225. https://doi.org/10.3389/fnbeh.2016.00225

Bose, M., Oliván, B., & Laferrère, B. (2009). Stress and obesity: the role of the hypothalamic-pituitary-adrenal axis in metabolic disease. Current Opinion in Endocrinology, Diabetes, and Obesity, 16(5), 340–346. https://doi.org/10.1097/MED.0b013e32832fa137

Boutelle, K. N., Wierenga, C. E., Bischoff-Grethe, A., Melrose, A. J., Grenesko-Stevens, E., Paulus, M. P., & Kaye, W. H. (2015). Increased brain response to appetitive tastes in the insula and amygdala in obese compared with healthy weight children when sated. International Journal of Obesity (2005), 39(4), 620–628. https://doi.org/10.1038/ijo.2014.206

Bunsey, M., & Eichenbaum, H. (1996). Conservation of hippocampal memory function in rats and humans. Nature, 379(6562), 255–257. https://doi.org/10.1038/379255a0

Burgos-Robles, A., Kimchi, E. Y., Izadmehr, E. M., Porzenheim, M. J., Ramos-Guasp, W. A., Nieh, E. H., Felix-Ortiz, A. C., Namburi, P., Leppla, C. A., Presbrey, K. N., Anandalingam, K. K., Pagan-Rivera, P. A., Anahtar, M., Beyeler, A., & Tye, K. M. (2017). Amygdala inputs to prefrontal cortex guide behavior amid conflicting cues of reward and punishment. Nature Neuroscience, 20(6), 824–835. https://doi.org/10.1038/nn.4553

Cohen, S. J., & Stackman, R. W. (2015). Assessing rodent hippocampal involvement in the novel object recognition task. A review. Behavioural Brain Research, 285, 105–117. https://doi.org/10.1016/j.bbr.2014.08.002

Connolly, L., Coveleskie, K., Kilpatrick, L. A., Labus, J. S., Ebrat, B., Stains, J., Jiang, Z., Tillisch, K., Raybould, H. E., & Mayer, E. A. (2013). Differences in brain responses between lean and obese women to a sweetened drink. Neurogastroenterology and Motility: The Official Journal of the European Gastrointestinal Motility Society, 25(7), 579–e460. https://doi.org/10.1111/nmo.12125

Cordner, Z. A., & Tamashiro, K. L. K. (2015). Effects of high-fat diet exposure on learning & memory. Physiology & Behavior, 152(Pt B), 363–371. https://doi.org/10.1016/j.physbeh.2015.06.008

de Andrade, A. M., Fernandes, M. da C., de Fraga, L. S., Porawski, M., Giovenardi, M., & Guedes, R. P. (2017). Omega-3 fatty acids revert high-fat diet-induced neuroinflammation but not recognition memory impairment in rats. Metabolic Brain Disease, 32(6), 1871–1881. https://doi.org/10.1007/s11011-017-0080-7

Décarie-Spain, L., Hryhorczuk, C., & Fulton, S. (2016). Dopamine signalling adaptations by prolonged high-fat feeding. Current Opinion in Behavioral Sciences, 9, 136–143. https://doi.org/10.1016/j.cobeha.2016.03.010

Del Olmo, N., & Ruiz-Gayo, M. (2018). Influence of High-Fat Diets Consumed During the Juvenile Period on Hippocampal Morphology and Function. Frontiers in Cellular Neuroscience, 12, 439. https://doi.org/10.3389/fncel.2018.00439

Elzinga, B. M., & Bremner, J. D. (2002). Are the neural substrates of memory the final common pathway in posttraumatic stress disorder (PTSD)? Journal of Affective Disorders, 70(1), 1–17. https://doi.org/10.1016/s0165-0327(01)00351-2

Ennaceur, A. (2010). One-trial object recognition in rats and mice: methodological and theoretical issues. Behavioural Brain Research, 215(2), 244–254. https://doi.org/10.1016/j.bbr.2009.12.036

Ennaceur, A., & Delacour, J. (1988). A new one-trial test for neurobiological studies of memory in rats. 1: Behavioral data. Behavioural Brain Research, 31(1), 47–59. https://doi.org/10.1016/0166-4328(88)90157-x

Felix-Ortiz, A. C., Beyeler, A., Seo, C., Leppla, C. A., Wildes, C. P., & Tye, K. M. (2013). BLA to vHPC inputs modulate anxiety-related behaviors. Neuron, 79(4), 658–664. https://doi.org/10.1016/j.neuron.2013.06.016

Francis, H., & Stevenson, R. (2013). The longer-term impacts of Western diet on human cognition and the brain. Appetite, 63, 119–128. https://doi.org/10.1016/j.appet.2012.12.018

Gomez, J. L., Bonaventura, J., Lesniak, W., Mathews, W. B., Sysa-Shah, P., Rodriguez, L. A., Ellis, R. J., Richie, C. T., Harvey, B. K., Dannals, R. F., Pomper, M. G., Bonci, A., & Michaelides, M. (2017). Chemogenetics revealed: DREADD occupancy and activation via converted clozapine. Science (New York, N.Y.), 357(6350), 503–507. https://doi.org/10.1126/science.aan2475

Hartley, T., Lever, C., Burgess, N., & O’Keefe, J. (2014). Space in the brain: how the hippocampal formation supports spatial cognition. Philosophical Transactions of the Royal Society of London. Series B, Biological Sciences, 369(1635), 20120510. https://doi.org/10.1098/rstb.2012.0510

Head, G. A. (2015). Cardiovascular and metabolic consequences of obesity. Frontiers in Physiology, 6, 32. https://doi.org/10.3389/fphys.2015.00032

Heyward, F. D., Gilliam, D., Coleman, M. A., Gavin, C. F., Wang, J., Kaas, G., Trieu, R., Lewis, J., Moulden, J., & Sweatt, J. D. (2016). Obesity Weighs down Memory through a Mechanism Involving the Neuroepigenetic Dysregulation of Sirt1. The Journal of Neuroscience: The Official Journal of the Society for Neuroscience, 36(4), 1324–1335. https://doi.org/10.1523/JNEUROSCI.1934-15.2016

Heyward, F. D., Walton, R. G., Carle, M. S., Coleman, M. A., Garvey, W. T., & Sweatt, J. D. (2012). Adult mice maintained on a high-fat diet exhibit object location memory deficits and reduced hippocampal SIRT1 gene expression. Neurobiology of Learning and Memory, 98(1), 25–32. https://doi.org/10.1016/j.nlm.2012.04.005

Hsu, T. M., Noble, E. E., Liu, C. M., Cortella, A. M., Konanur, V. R., Suarez, A. N., Reiner, D. J., Hahn, J. D., Hayes, M. R., & Kanoski, S. E. (2018). A hippocampus to prefrontal cortex neural pathway inhibits food motivation through glucagon-like peptide-1 signaling. Molecular Psychiatry, 23(7), 1555–1565. https://doi.org/10.1038/mp.2017.91

Ilg, A.-K., Enkel, T., Bartsch, D., & Bähner, F. (2018). Behavioral Effects of Acute Systemic Low-Dose Clozapine in Wild-Type Rats: Implications for the Use of DREADDs in Behavioral Neuroscience. Frontiers in Behavioral Neuroscience, 12, 173. https://doi.org/10.3389/fnbeh.2018.00173

Kaouane, N., Porte, Y., Vallée, M., Brayda-Bruno, L., Mons, N., Calandreau, L., Marighetto, A., Piazza, P. V., & Desmedt, A. (2012). Glucocorticoids can induce PTSD-like memory impairments in mice. Science (New York, N.Y.), 335(6075), 1510–1513. https://doi.org/10.1126/science.1207615

Kendig, M. D., Westbrook, R. F., & Morris, M. J. (2019). Pattern of access to cafeteria-style diet determines fat mass and degree of spatial memory impairments in rats. Scientific Reports, 9(1), 13516. https://doi.org/10.1038/s41598-019-50113-3

Khan, N. A., Baym, C. L., Monti, J. M., Raine, L. B., Drollette, E. S., Scudder, M. R., Moore, R. D., Kramer, A. F., Hillman, C. H., & Cohen, N. J. (2015). Central adiposity is negatively associated with hippocampal-dependent relational memory among overweight and obese children. The Journal of Pediatrics, 166(2), 302–308.e1. https://doi.org/10.1016/j.jpeds.2014.10.008

Khazen, T., Hatoum, O. A., Ferreira, G., & Maroun, M. (2019). Acute exposure to a high-fat diet in juvenile male rats disrupts hippocampal-dependent memory and plasticity through glucocorticoids. Scientific Reports, 9(1), 12270. https://doi.org/10.1038/s41598-019-48800-2

Kim, J. M., Kim, D. H., Lee, Y., Park, S. J., & Ryu, J. H. (2014). Distinct roles of the hippocampus and perirhinal cortex in GABAA receptor blockade-induced enhancement of object recognition memory. Brain Research, 1552, 17–25. https://doi.org/10.1016/j.brainres.2014.01.024

Kosari, S., Badoer, E., Nguyen, J. C. D., Killcross, A. S., & Jenkins, T. A. (2012). Effect of western and high fat diets on memory and cholinergic measures in the rat. Behavioural Brain Research, 235(1), 98–103. https://doi.org/10.1016/j.bbr.2012.07.017

Lavin, D. N., Joesting, J. J., Chiu, G. S., Moon, M. L., Meng, J., Dilger, R. N., & Freund, G. G. (2011). Fasting induces an anti-inflammatory effect on the neuroimmune system which a high-fat diet prevents. Obesity (Silver Spring, Md.), 19(8), 1586–1594. https://doi.org/10.1038/oby.2011.73

Layton, B., & Krikorian, R. (2002). Memory mechanisms in posttraumatic stress disorder. The Journal of Neuropsychiatry and Clinical Neurosciences, 14(3), 254–261. https://doi.org/10.1176/jnp.14.3.254

LeDoux, J. (2003). The emotional brain, fear, and the amygdala. Cellular and Molecular Neurobiology, 23(4–5), 727–738. https://doi.org/10.1023/a:1025048802629

Llorca-Torralba, M., Suárez-Pereira, I., Bravo, L., Camarena-Delgado, C., Garcia-Partida, J. A., Mico, J. A., & Berrocoso, E. (2019). Chemogenetic Silencing of the Locus Coeruleus-Basolateral Amygdala Pathway Abolishes Pain-Induced Anxiety and Enhanced Aversive Learning in Rats. Biological Psychiatry, 85(12), 1021–1035. https://doi.org/10.1016/j.biopsych.2019.02.018

MacLaren, D. A. A., Browne, R. W., Shaw, J. K., Krishnan Radhakrishnan, S., Khare, P., España, R. A., & Clark, S. D. (2016). Clozapine N-Oxide Administration Produces Behavioral Effects in Long-Evans Rats: Implications for Designing DREADD Experiments. ENeuro, 3(5). https://doi.org/10.1523/ENEURO.0219-16.2016

Mahan, A. L., & Ressler, K. J. (2012). Fear conditioning, synaptic plasticity and the amygdala: implications for posttraumatic stress disorder. Trends in Neurosciences, 35(1), 24–35. https://doi.org/10.1016/j.tins.2011.06.007

Malnick, S. D. H., & Knobler, H. (2006). The medical complications of obesity. QJM: Monthly Journal of the Association of Physicians, 99(9), 565–579. https://doi.org/10.1093/qjmed/hcl085

Mansur, R. B., Brietzke, E., & McIntyre, R. S. (2015). Is there a “metabolic-mood syndrome”? A review of the relationship between obesity and mood disorders. Neuroscience and Biobehavioral Reviews, 52, 89–104. https://doi.org/10.1016/j.neubiorev.2014.12.017

Maroun, M., & Akirav, I. (2008). Arousal and stress effects on consolidation and reconsolidation of recognition memory. Neuropsychopharmacology: Official Publication of the American College of Neuropsychopharmacology, 33(2), 394–405. https://doi.org/10.1038/sj.npp.1301401

Martin, A. A., & Davidson, T. L. (2014). Human cognitive function and the obesogenic environment. Physiology & Behavior, 136, 185–193. https://doi.org/10.1016/j.physbeh.2014.02.062

McCormick, C. M., & Mathews, I. Z. (2010). Adolescent development, hypothalamic-pituitary-adrenal function, and programming of adult learning and memory. Progress in Neuro-Psychopharmacology & Biological Psychiatry, 34(5), 756–765. https://doi.org/10.1016/j.pnpbp.2009.09.019

McEwen, B. S., Nasca, C., & Gray, J. D. (2016). Stress Effects on Neuronal Structure: Hippocampus, Amygdala, and Prefrontal Cortex. Neuropsychopharmacology: Official Publication of the American College of Neuropsychopharmacology, 41(1), 3–23. https://doi.org/10.1038/npp.2015.171

McGaugh, J. L. (2004). The amygdala modulates the consolidation of memories of emotionally arousing experiences. Annual Review of Neuroscience, 27, 1–28. https://doi.org/10.1146/annurev.neuro.27.070203.144157

McLean, F. H., Grant, C., Morris, A. C., Horgan, G. W., Polanski, A. J., Allan, K., Campbell, F. M., Langston, R. F., & Williams, L. M. (2018). Rapid and reversible impairment of episodic memory by a high-fat diet in mice. Scientific Reports, 8(1), 11976. https://doi.org/10.1038/s41598-018-30265-4

Mestre, Z. L., Bischoff-Grethe, A., Eichen, D. M., Wierenga, C. E., Strong, D., & Boutelle, K. N. (2017). Hippocampal atrophy and altered brain responses to pleasant tastes among obese compared with healthy weight children. International Journal of Obesity (2005), 41(10), 1496–1502. https://doi.org/10.1038/ijo.2017.130

Morin, J.-P., Rodríguez-Durán, L. F., Guzmán-Ramos, K., Perez-Cruz, C., Ferreira, G., Diaz-Cintra, S., & Pacheco-López, G. (2017). Palatable Hyper-Caloric Foods Impact on Neuronal Plasticity. Frontiers in Behavioral Neuroscience, 11, 19. https://doi.org/10.3389/fnbeh.2017.00019

Mucellini, A. B., Laureano, D. P., Silveira, P. P., & Sanvitto, G. L. (2019). Maternal and post-natal obesity alters long-term memory and hippocampal molecular signaling of male rat. Brain Research, 1708, 138–145. https://doi.org/10.1016/j.brainres.2018.12.021

Mueller, K., Sacher, J., Arelin, K., Holiga, S., Kratzsch, J., Villringer, A., & Schroeter, M. L. (2012). Overweight and obesity are associated with neuronal injury in the human cerebellum and hippocampus in young adults: a combined MRI, serum marker and gene expression study. Translational Psychiatry, 2, e200. https://doi.org/10.1038/tp.2012.121

Murray, S., & Chen, E. Y. (2019). Examining Adolescence as a Sensitive Period for High-Fat, High-Sugar Diet Exposure: A Systematic Review of the Animal Literature. Frontiers in Neuroscience, 13, 1108. https://doi.org/10.3389/fnins.2019.01108

Mutlu, O., Ulak, G., Celikyurt, I. K., Tanyeri, P., Akar, F. Y., & Erden, F. (2011). Effects of olanzapine and clozapine on memory acquisition, consolidation and retrieval in mice using the elevated plus maze test. Neuroscience Letters, 501(3), 143–147. https://doi.org/10.1016/j.neulet.2011.07.004

Naneix, F., Tantot, F., Glangetas, C., Kaufling, J., Janthakhin, Y., Boitard, C., De Smedt-Peyrusse, V., Pape, J. R., Vancassel, S., Trifilieff, P., Georges, F., Coutureau, E., & Ferreira, G. (2017). Impact of Early Consumption of High-Fat Diet on the Mesolimbic Dopaminergic System. ENeuro, 4(3). https://doi.org/10.1523/ENEURO.0120-17.2017

Noble, E. E., & Kanoski, S. E. (2016). Early life exposure to obesogenic diets and learning and memory dysfunction. Current Opinion in Behavioral Sciences, 9, 7–14. https://doi.org/10.1016/j.cobeha.2015.11.014

Nyaradi, A., Foster, J. K., Hickling, S., Li, J., Ambrosini, G. L., Jacques, A., & Oddy, W. H. (2014). Prospective associations between dietary patterns and cognitive performance during adolescence. Journal of Child Psychology and Psychiatry, and Allied Disciplines, 55(9), 1017–1024. https://doi.org/10.1111/jcpp.12209

Ogden, C. L., Carroll, M. D., Lawman, H. G., Fryar, C. D., Kruszon-Moran, D., Kit, B. K., & Flegal, K. M. (2016). Trends in Obesity Prevalence Among Children and Adolescents in the United States, 1988-1994 Through 2013-2014. JAMA, 315(21), 2292–2299. https://doi.org/10.1001/jama.2016.6361

Okuda, S., Roozendaal, B., & McGaugh, J. L. (2004). Glucocorticoid effects on object recognition memory require training-associated emotional arousal. Proceedings of the National Academy of Sciences of the United States of America, 101(3), 853–858. https://doi.org/10.1073/pnas.0307803100

Okuyama, T., Kitamura, T., Roy, D. S., Itohara, S., & Tonegawa, S. (2016). Ventral CA1 neurons store social memory. Science (New York, N.Y.), 353(6307), 1536–1541. https://doi.org/10.1126/science.aaf7003

Oliveira, A. M. M., Hawk, J. D., Abel, T., & Havekes, R. (2010). Post-training reversible inactivation of the hippocampus enhances novel object recognition memory. Learning & Memory (Cold Spring Harbor, N.Y.), 17(3), 155–160. https://doi.org/10.1101/lm.1625310

Øverby, N. C., Lüdemann, E., & Høigaard, R. (2013). Self-reported learning difficulties and dietary intake in Norwegian adolescents. Scandinavian Journal of Public Health, 41(7), 754–760. https://doi.org/10.1177/1403494813487449

Pagoto, S. L., Schneider, K. L., Bodenlos, J. S., Appelhans, B. M., Whited, M. C., Ma, Y., & Lemon, S. C. (2012). Association of post-traumatic stress disorder and obesity in a nationally representative sample. Obesity (Silver Spring, Md.), 20(1), 200–205. https://doi.org/10.1038/oby.2011.318

Paré, D. (2003). Role of the basolateral amygdala in memory consolidation. Progress in Neurobiology, 70(5), 409–420. https://doi.org/10.1016/s0301-0082(03)00104-7

Pasquali, R., Vicennati, V., Cacciari, M., & Pagotto, U. (2006). The hypothalamic-pituitary-adrenal axis activity in obesity and the metabolic syndrome. Annals of the New York Academy of Sciences, 1083, 111–128. https://doi.org/10.1196/annals.1367.009

Paxinos, G., & Watson, C. (2007). The Rat brain in Stereotaxic Coordinates (6th ed.). Elsevier.

Perkonigg, A., Owashi, T., Stein, M. B., Kirschbaum, C., & Wittchen, H.-U. (2009). Posttraumatic stress disorder and obesity: evidence for a risk association. American Journal of Preventive Medicine, 36(1), 1–8. https://doi.org/10.1016/j.amepre.2008.09.026

Phillips, M. L., Robinson, H. A., & Pozzo-Miller, L. (2019). Ventral hippocampal projections to the medial prefrontal cortex regulate social memory. ELife, 8. https://doi.org/10.7554/eLife.44182

Pitkänen, A., Pikkarainen, M., Nurminen, N., & Ylinen, A. (2000). Reciprocal connections between the amygdala and the hippocampal formation, perirhinal cortex, and postrhinal cortex in rat. A review. Annals of the New York Academy of Sciences, 911, 369–391. https://doi.org/10.1111/j.1749-6632.2000.tb06738.x

Rei, D., Mason, X., Seo, J., Gräff, J., Rudenko, A., Wang, J., Rueda, R., Siegert, S., Cho, S., Canter, R. G., Mungenast, A. E., Deisseroth, K., & Tsai, L.-H. (2015). Basolateral amygdala bidirectionally modulates stress-induced hippocampal learning and memory deficits through a p25/Cdk5-dependent pathway. Proceedings of the National Academy of Sciences of the United States of America, 112(23), 7291–7296. https://doi.org/10.1073/pnas.1415845112

Reichelt, A. C. (2016). Adolescent Maturational Transitions in the Prefrontal Cortex and Dopamine Signaling as a Risk Factor for the Development of Obesity and High Fat/High Sugar Diet Induced Cognitive Deficits. Frontiers in Behavioral Neuroscience, 10, 189. https://doi.org/10.3389/fnbeh.2016.00189

Reichelt, A. C., & Rank, M. M. (2017). The impact of junk foods on the adolescent brain. Birth Defects Research, 109(20), 1649–1658. https://doi.org/10.1002/bdr2.1173

Rogan, S. C., & Roth, B. L. (2011). Remote control of neuronal signaling. Pharmacological Reviews, 63(2), 291–315. https://doi.org/10.1124/pr.110.003020

Roozendaal, B., Castello, N. A., Vedana, G., Barsegyan, A., & McGaugh, J. L. (2008). Noradrenergic activation of the basolateral amygdala modulates consolidation of object recognition memory. Neurobiology of Learning and Memory, 90(3), 576–579. https://doi.org/10.1016/j.nlm.2008.06.010

Roozendaal, B., Okuda, S., Van der Zee, E. A., & McGaugh, J. L. (2006). Glucocorticoid enhancement of memory requires arousal-induced noradrenergic activation in the basolateral amygdala. Proceedings of the National Academy of Sciences of the United States of America, 103(17), 6741–6746. https://doi.org/10.1073/pnas.0601874103

Sahoo, K., Sahoo, B., Choudhury, A. K., Sofi, N. Y., Kumar, R., & Bhadoria, A. S. (2015). Childhood obesity: causes and consequences. Journal of Family Medicine and Primary Care, 4(2), 187–192. https://doi.org/10.4103/2249-4863.154628

Saygin, Z. M., Osher, D. E., Koldewyn, K., Martin, R. E., Finn, A., Saxe, R., Gabrieli, J. D. E., & Sheridan, M. (2015). Structural connectivity of the developing human amygdala. PloS One, 10(4), e0125170. https://doi.org/10.1371/journal.pone.0125170

Sellbom, K. S., & Gunstad, J. (2012). Cognitive function and decline in obesity. Journal of Alzheimer’s Disease: JAD, 30 Suppl 2, S89–95. https://doi.org/10.3233/JAD-2011-111073

Spear, L. P. (2000). The adolescent brain and age-related behavioral manifestations. Neuroscience and Biobehavioral Reviews, 24(4), 417–463. https://doi.org/10.1016/s0149-7634(00)00014-2

Stuber, G. D., Sparta, D. R., Stamatakis, A. M., van Leeuwen, W. A., Hardjoprajitno, J. E., Cho, S., Tye, K. M., Kempadoo, K. A., Zhang, F., Deisseroth, K., & Bonci, A. (2011). Excitatory transmission from the amygdala to nucleus accumbens facilitates reward seeking. Nature, 475(7356), 377–380. https://doi.org/10.1038/nature10194

Tantot, F., Parkes, S. L., Marchand, A. R., Boitard, C., Naneix, F., Layé, S., Trifilieff, P., Coutureau, E., & Ferreira, G. (2017). The effect of high-fat diet consumption on appetitive instrumental behavior in rats. Appetite, 108, 203–211. https://doi.org/10.1016/j.appet.2016.10.001

Tran, D. M. D., & Westbrook, R. F. (2015). Rats Fed a Diet Rich in Fats and Sugars Are Impaired in the Use of Spatial Geometry. Psychological Science, 26(12), 1947–1957. https://doi.org/10.1177/0956797615608240

Tran, D. M. D., & Westbrook, R. F. (2017). A high-fat high-sugar diet-induced impairment in place-recognition memory is reversible and training-dependent. Appetite, 110, 61–71. https://doi.org/10.1016/j.appet.2016.12.010

Tran, D. M. D., & Westbrook, R. F. (2018). Dietary effects on object recognition: The impact of high-fat high-sugar diets on recollection and familiarity-based memory. Journal of Experimental Psychology. Animal Learning and Cognition, 44(3), 217–228. https://doi.org/10.1037/xan0000170

Tucker, K. R., Godbey, S. J., Thiebaud, N., & Fadool, D. A. (2012). Olfactory ability and object memory in three mouse models of varying body weight, metabolic hormones, and adiposity. Physiology & Behavior, 107(3), 424–432. https://doi.org/10.1016/j.physbeh.2012.09.007

van Galen, K. A., Ter Horst, K. W., Booij, J., la Fleur, S. E., & Serlie, M. J. (2018). The role of central dopamine and serotonin in human obesity: lessons learned from molecular neuroimaging studies. Metabolism: Clinical and Experimental, 85, 325–339. https://doi.org/10.1016/j.metabol.2017.09.007

Walls, H. L., Backholer, K., Proietto, J., & McNeil, J. J. (2012). Obesity and trends in life expectancy. Journal of Obesity, 2012, 107989. https://doi.org/10.1155/2012/107989

Wang, C., Chan, J. S. Y., Ren, L., & Yan, J. H. (2016). Obesity Reduces Cognitive and Motor Functions across the Lifespan. Neural Plasticity, 2016, 2473081. https://doi.org/10.1155/2016/2473081

Widya, R. L., de Roos, A., Trompet, S., de Craen, A. J., Westendorp, R. G., Smit, J. W., van Buchem, M. A., van der Grond, J., & PROSPER Study Group. (2011). Increased amygdalar and hippocampal volumes in elderly obese individuals with or at risk of cardiovascular disease. The American Journal of Clinical Nutrition, 93(6), 1190–1195. https://doi.org/10.3945/ajcn.110.006304

Yeomans, M. R. (2017). Adverse effects of consuming high fat-sugar diets on cognition: implications for understanding obesity. The Proceedings of the Nutrition Society, 76(4), 455–465. https://doi.org/10.1017/S0029665117000805

Zuloaga, K. L., Johnson, L. A., Roese, N. E., Marzulla, T., Zhang, W., Nie, X., Alkayed, F. N., Hong, C., Grafe, M. R., Pike, M. M., Raber, J., & Alkayed, N. J. (2016). High fat diet-induced diabetes in mice exacerbates cognitive deficit due to chronic hypoperfusion. Journal of Cerebral Blood Flow and Metabolism: Official Journal of the International Society of Cerebral Blood Flow and Metabolism, 36(7), 1257–1270. https://doi.org/10.1177/0271678X15616400

